# Artificial upward trends in Greek marine landings: a case of presentist bias in European fisheries

**DOI:** 10.1101/859694

**Authors:** Athanassios C. Tsikliras, Donna Dimarchopoulou, Androniki Pardalou

## Abstract

According to the official landings as reported by the international databases for Greece, the declining trend of the Greek marine fisheries landings that had been continuous since the mid 1990s has been reversed during the last two years, with the total marine fisheries landings showing elevated catches after 2016. We claim that this upward trend is an artifact that is attributed to the combined reporting of the landings of additional fleets since 2016 that had been separately reported before and resulted in 20-30% inflation of the landings. In 2016, the Greek statistical authorities included the landings of 10 000 small-scale coastal vessels with engine horsepower lower than 20 HP together with the remaining coastal vessels, purse-seiners and trawlers whose landings formed the official reported Greek marine fisheries landings from 1970 to 2015. We acknowledge that this act of partial catch reconstruction improved the resolution of the landings and the officially reported values are now more realistic. However, the artificial, albeit inadvertent, inflation of the official Greek marine fisheries landings as they appear in international databases is a clear case of ‘presentist bias’ and may distort stock assessments and ecosystem modeling. As the currently misleading data stand, they are cause for substantial misinterpretation and analytical errors that can influence fisheries policy and have serious implications for fisheries management. We suggest that researchers should refrain from using the combined time-series and that a correction should be applied to the original time series (1970-2015) to account for the entire small-scale coastal fleet.

## 1. Introduction

The recent reconstruction of global catches revealed that a large proportion of the total official catch statistics still remains unreported and that unreported catch mainly refers to small-scale artisanal and recreational fisheries, which are incompletely recorded or even omitted from official catch statistics on a global scale [1]. The underreporting or misreporting of marine catches and fishing effort may bias stock assessments and hamper fisheries management because a large proportion of the total biomass removed by fishing is not accounted for [1, 2]. This, in turn, may influence national policies on fisheries [3].

In Greek waters, a fisheries data-poor area [4], the initial [5] and recent catch reconstructions [6] revealed several discrepancies, inconsistencies and dataset discontinuities prior to 1982 and an unreported catch that may exceed 30% of the officially reported landings. However, despite those inconsistencies and biases that are confined to the earlier years of the dataset (1970-1981) and several aggregations to higher taxonomic groups [7], the *modus operandi* of the Greek statistical authorities remained the same between 1982 and 2015 [8, 9]. From 2016 onwards, the Greek statistical authorities reformed their reporting by changing the number of species that are being recorded and by including the landings derived from small scale coastal vessels with engine lower than 20 HP (around 10 000 vessels) to the landings of the remaining coastal vessels, purse-seiners and trawlers whose landings had formed the official reported Greek marine fisheries statistics from 1970 to 2015 [10] to the General Fisheries Commission for the Mediterranean (GFCM) and Food and Agricultural Organization (FAO) databases. This change that improved the reliability of reporting, is clearly stated in the 2016 Annual Bulletin [11], which is available online: *“Until the reference year 2015, the survey covered the professional motor-propelled fishing vessels of 20 HP and over. From the reference year 2016 onwards, the survey covers all motor-propelled professional fishing vessels irrespective of their horsepower”*. This statement, however, does not appear in internationa databases of GFCM and FAO and constitutes a clear case of ‘presentist bias’, a term that has been recently proposed in fisheries science to describe the overestimation of the present against the past [12]. It refers to an improvement in an official catch reporting system (such as an inclusion of a previously unmonitored activity related to fleet, gear or region) that leads to an increase in recent reported catches without the corresponding past (unmonitored) catches being corrected for retroactively [3]. ‘Presentist bias’ is an inadvertent consequence of the effort most countries make to improve their national data collection and reporting systems [3].

The objective of the present work was to point out the first case of ‘presentist bias’ in European fisheries that emerged from including the landings of additional fleets to the official fisheries statistics without accounting for the associated fishing effort increase, or without correcting these data backwards (hindcasting) to generate a full historical time series. Such incomplete time series have serious implications for future stock assessments and policy development because they result in misinterpretation of fisheries trends, as well as on ecosystem modeling and studies of climate impact on marine populations.

## 2. Materials and methods

The annual catches of the Greek marine fishing fleet, expressed as live weight equivalent of landings, had been recorded for 66 species or groups of species (henceforth called taxa) since 1982 by the Greek statistical authorities (the current official name of the service is Hellenic Statistical Authority, HELSTAT) and published in annual bulletins (www.statistics.gr/en/). The HELSTAT dataset originally referred to the legal and reported offshore (trawlers and purse-seiners) and coastal fisheries (netters, boat seiners and longliners with engine horsepower greater than 20 HP) landings, excluding discarded, illegal and unreported catches, recreational and sport fishing as well as the catch of small scale coastal vessels with engine horsepower lower than 20 HP [5]. Although the latter was partly recorded by another national statistical authority and was available only in Greek, it was not a part of the official Greek marine fisheries landings up to 2015.

Up to 2015 the landings of four fleets (trawlers, purse seiners, boat seiners, small-scale coastal vessels with engine horsepower > 20 HP) are reported annually and their landings were acquired from the official HELLSTAT annual bulletins since 1982. From 2016, onwards the landings of a fifth fleet (small-scale coastal vessels with engine horsepower < 20 HP) were added in the official Greek marine landings dataset and these data were also acquired from the official HELLSTAT annual bulletins. In this work, the years 2016 and 2017 were spilt into landings of four and five fleets for presentation purposes.

GFCM records the annual landings of the Mediterranean and Black Sea fisheries stocks since 1970 on a country, species and subdivision basis whereas the FAO dataset begins in 1950 and differs from that of GFCM in terms of spatial division and species aggregation. Both FAO and GFCM catch statistics are available online and can be downloaded using the FishStat J software [13]. The equivalent to the HELLSTAT dataset landings for Greece were downloaded from both FAO and GFCM databases for the same period of time. It should be noted here that the GFCM/FAO only include the landings of the HELLSTAT, without any information on fleets and effort data [13].

The inclusion of all fleets in official Greek marine fisheries landings statistics is essential and a main target of catch reconstructions [1, 3]. For this reason, a correction of including the landings of a fifth fleet (small-scale coastal vessels with engine horsepower < 20 HP) was applied to the entire dataset (1982-2017) here, following the catch reconstruction methodology [5, 6]. Finally, the catch per unit of effort (CPUE) was estimated as landings (in metric tons) per vessel for the four and the five fleets aiming to reveal the impact of adding the landings of an extra fleet of 10 000 small-scale coastal fishing vessels to the official marine fisheries landings time series.

## 3. Results

The official Greek marine fisheries landings derived from the four fleets (trawlers, purse-seiners, boat-seiners and small-scale coastal vessels with engine horsepower >20HP), as they are reported by HELSTAT and adopted by GFCM and FAO datasets have been increasing up to the mid 1990s and declining from their historical maximum in 1994 (165 000 t) to 2012 when they stabilized around 60 000 t (blue line in Figure 1). The landings of the four fleets continue to fluctuate around 60 000 t in 2016 and 2017 (dotted blue line in Figure 1).

**Figure 1.**
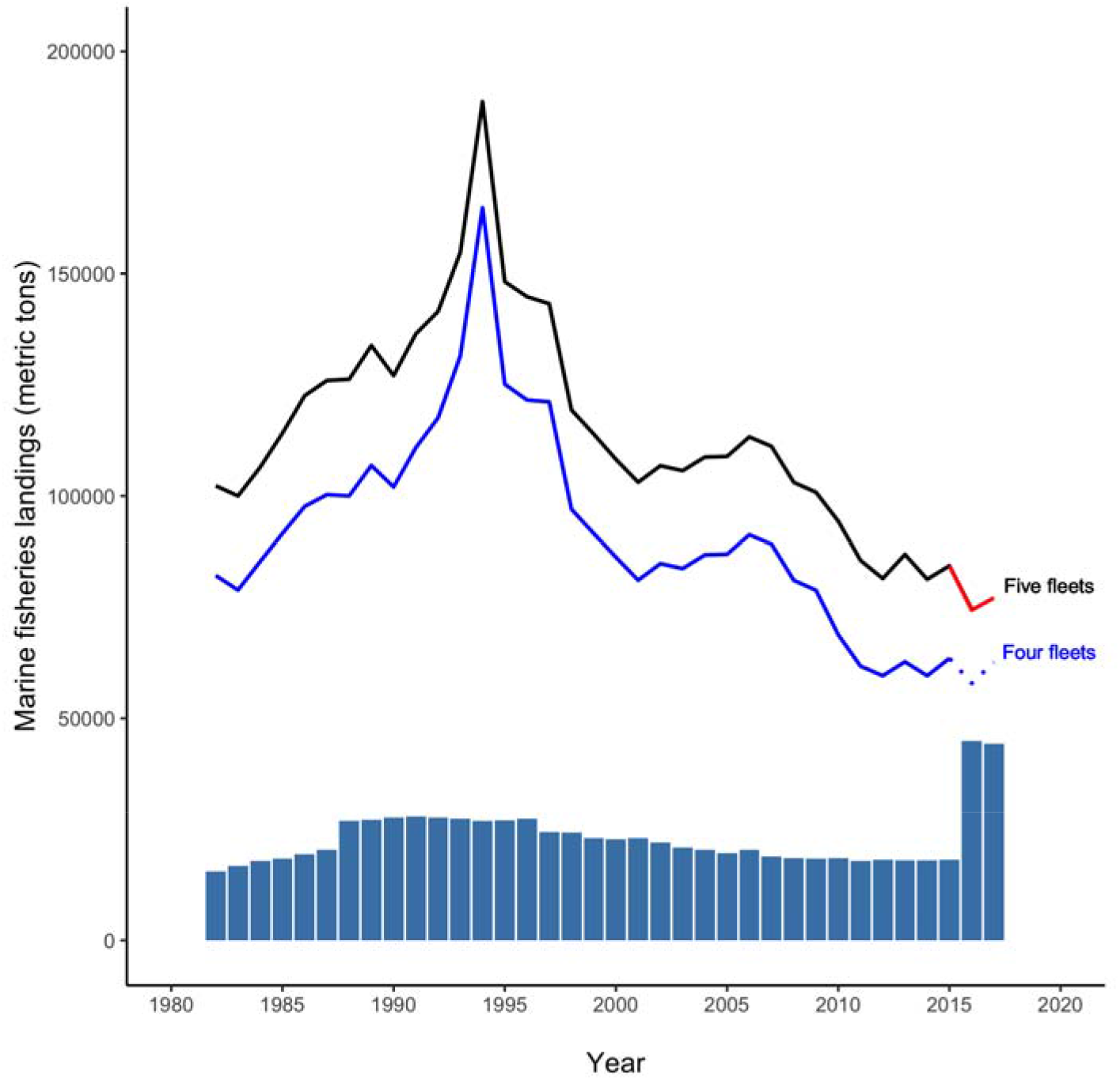
The official Greek marine fisheries landings without (blue line, four fleets) and with (black line, five fleets) the small-scale coastal vessels with engine horsepower <20 HP for 2016 and 2017 (red line) and hindcasted for 1982-2015 based on the recent catch reconstruction of Greek marine fisheries landings [5, 6]. The bars indicate the annual number of vessels from which the landings are derived.

The inclusion of the landings of the around 10 000 vessels forming a fifth fleet (small-scale coastal vessels with engine horsepower <20HP) in 2016, inflated the official Greek marine fisheries landings in 2016 and 2017 by 28 and 23%, respectively (red line in Figure 1). The hindcast of the five fleets follows the same fluctuation with that of the four fleets but the magnitude is higher (black line in Figure 1). The landings in the FAO and GFCM datasets are slightly different in the earlier years and identical in the latest years compared to the HELLSTAT dataset but showing the exact same trend up to 2015 and follow the five fleet dataset from 2016 onwards.

The artificial inflation of including an extra fleet in the dataset also caused the abrupt dropping of the overall CPUE to half of its previous values, from 10.5 t/vessel to around 5 t/vessel and resulted in a steeper decline of the CPUE of the Greek marine fisheries across the study period (Figure 2).

**Figure 2.**
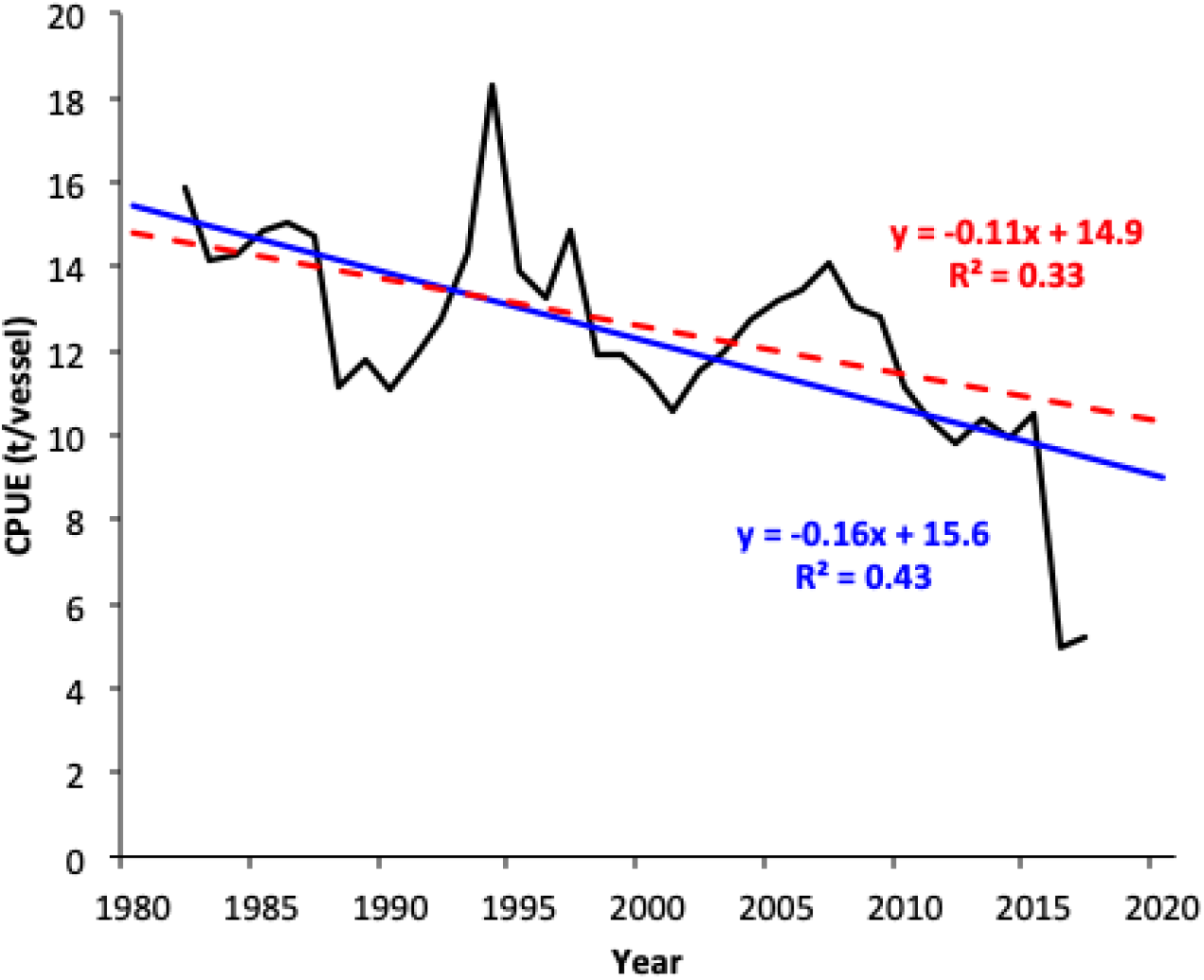
The CPUE (tons of landings per vessel) based on the official Greek marine fisheries landings. The blue trend line refers to the mixed dataset (four fleets for 1982-2015, five fleets for 2016-2017) whereas the red dotted line refers to the corrected dataset (five fleets across the study period, reconstructed landings for 1982-2015, original landings for 2016-2017).

## 4. Discussion

The official Greek marine fisheries landings have been extensively used to assess the status of many stocks in the eastern Mediterranean Sea using catch-based methods [8, 10] and surplus production models [14], in several ecosystem models in the area [15], in examining the effect of sea warming [16] and climate variability [17] on marine populations and the effect of fisheries on marine ecosystems [10].

The partial reconstruction by the inclusion of the landings of 10 000 small-scale coastal vessels is certainly an improvement of the monitoring system and bridges the gap between official statistics and the actual biomass of marine organisms removed by fishing. Therefore, it is acknowledged as a very positive step but it is also the first documented case of presentist bias in European fisheries. The transition from a recording system of four fleets to a recording system of five fleets results in an artificial inflation of reported landings, which will certainly affect the scientific output of various disciplines and distort fisheries management [3]. Firstly, a real 20-30% increase in landings would suggest that the fisheries regulations and the general management scheme in the Greek seas had been successful. However, all recent publications and assessments [10, 14, 18, 19] highlight the bad status of the Greek stocks and the need for less fishing, a condition that holds for most Mediterranean fisheries [10, 14]. This clearly opposes the increased fishing pressure applied to pelagic and demersal stocks as a result of “fishing in international waters” of the Greek fleet that practically allows them to operate all year round [20]. It has been recently reported that all species targeted by the fishing fleets are being overexploited in the Mediterranean Sea [21], with none of the Mediterranean stocks being sustainably exploited [22], including 98% of unassessed demersal stocks [23]. Secondly, the official landings are being used in stock assessments and are particularly important in catch-based method for estimating stock status [24] and surplus production models for estimating fisheries reference points [25]. An artificial change of the catch in the last 2-3 years of the time-series will affect the reference points and the output of the stock status [26] and will influence national policy discussions on fisheries and fisheries management decisions [3]. Thirdly, the inclusion of the landings of a coastal fishing fleet disproportionally affects the landings composition in favour of species caught by netters and long-liners. This in turn distorts the input to ecosystem models and to methods that rely on landings and are used to determine the effect of fishing (such as mean weighted trophic level of the catch: [27]) and climate (such as the mean temperature of the catch: [28]) on marine populations and ecosystems. The total Greek marine fisheries landings can be rather easily updated to include the landings of the fifth fleet back to 1982 (black line in Figure 1) because the data are available in the recent catch reconstruction [5, 6]. However, the landings of individual taxa, which are used in stock assessments and ecosystem models, are harder to obtain. In any case, reforming of a landings time-series by including an additional fleet of 10 000 coastal vessels requires that all users, especially international ones that use the GFCM/FAO datasets, should be clearly informed and given the opportunity to convert the two time series so they have the same fleets throughout the dataset. We by no means imply that HELLSTAT and GFCM/FAO should be blamed for doing something wrong or are deliberately misreporting catch statistics but this artificial inflation is an issue they, especially GFCM/FAO, should address.

In conclusion, we acknowledge that this act of partial reconstruction improved the resolution of the landings and is more realistic and close to the actual biomass removed from the sea, but the inflation of landings in international datasets has serious implications for stock assessments, ecosystem models and climate impact studies. We propose correction of the original time series (from at least 1982 to 2015) to account for the entire small-scale coastal fleet based on the recent reconstruction of marine fisheries landings of Greece [5, 6], which already exists. GFCM/FAO should be urgently informed to correct the international records. Finally, we suggest users to refrain from mixing the two datasets because they will get unrealistic, even erroneous, results.

